# Unraveling the critical growth factors for stable cultivation of (nano-sized) Micrarchaeota

**DOI:** 10.1101/2021.04.28.441856

**Authors:** Susanne Krause, Sabrina Gfrerer, Carsten Reuse, Nina Dombrowski, Laura Villanueva, Boyke Bunk, Cathrin Spröer, Thomas R. Neu, Ute Kuhlicke, Kerstin Schmidt-Hohagen, Karsten Hiller, Reinhard Rachel, Anja Spang, Johannes Gescher

**Affiliations:** Department of Applied Biology, Karlsruhe Institute of Technology (KIT), Karlsruhe, Germany; Institute for Biological Interfaces, Karlsruhe Institute of Technology (KIT), Eggenstein-Leopoldshafen, Germany; Department of Marine Microbiology and Biogeochemistry, NIOZ, Royal Netherlands Institute for Sea Research, Den Burg, The Netherlands; Department of Earth Sciences, Faculty of Geosciences, Utrecht University, Utrecht, The Netherlands; Leibniz Institute DSMZ, Braunschweig, Germany; Helmholtz-Centre for Environmental Research UFZ, Magdeburg, Germany; Bioinformatics & Biochemistry, Technische Universität Braunschweig, Braunschweig, Germany; Braunschweig Integrated Centre for Systems Biology (BRICS), Technische Universität Braunschweig, Braunschweig, Germany; Center for Electron Microscopy, University of Regensburg, Regensburg, Germany; Department of Cell- and Molecular Biology, Science for Life Laboratory, Uppsala University, SE-75123, Uppsala, Sweden

**Author notes:** Corresponding author: Prof. Dr. Johannes Gescher, Karlsruhe Institute of Technology, Institute of Applied Biosciences, Department of applied biology. These authors contributed equally to this work (Susanne Krause, Sabrina Gfrerer).

## Abstract

Micrarchaeota are members of the archaeal DPANN superphylum. These so far poorly characterized archaea have been found to have reduced genomes and likely depend on interactions with host organisms for growth and survival. Here we report on the enrichment of the first stable co-culture of a member of the Micrarchaeota together with its host, as well as the isolation of the latter. Electron microscopic analysis suggest that growth is dependent on the physical interaction of the two organisms within a biofilm. The interaction seems to be ensured by the necessity to grow in form of a biofilm. Furthermore, transcriptomic analyses indicate a shift towards biofilm formation of the host as a result of co-cultivation. Finally, genomic, metabolomic, extracellular polymeric substance (EPSs) and lipid content analyses reveal that the Micrarchaeon symbiont relies on the acquisition of metabolites from its host and thereby provide first insights into the basis of symbiont-host interactions.

## Introduction

In 2002, Huber and colleagues described a novel nano-sized archaeon, *Nanoarchaeum equitans* ^1^. Later, metagenomic data of environmental samples revealed that the Nanoarchaeota are part of a tentative superphylum of nano-sized archaea now referred to as DPANN – an acronym on its first members lineages, the Diapherotrites, Parvarchaeota, Aenigmarchaeota, Nanoarchaeota, and Nanohaloarchaeota ^2^. Most DPANN representatives have reduced genomes and are thought to comprise a diversity of potential archaeal symbionts. Besides the name-giving phyla, the DPANN also include the Woese- and Pacearchaeota ^3^, Huberarchaeota ^4^, Micrarchaeota ^5^, Altiarchaeota ^6^, Undinarchaeota ^7^ and Mamarchaeota ^8,9^ as well as several so far undefined phyla ^8,10^. Nano-sized archaea are globally distributed and can comprise non-negligible proportions of microbial communities ^11^. Yet only a few representatives have been enriched under laboratory conditions ^1,12–17^. Current genomic data suggest that most DPANN archaea have reduced genomes, limited metabolic capabilities and various auxotrophies and might depend on interactions with other community members. The extent of genome reduction varies within the DPANN members. For instance, marine Nanoarchaeota are characterized by highly reduced genomes of about 0.5 Mbp and seem to represent ectoparasites that are strongly dependent on their host ^1^. On the other hand, the first members of the Nanohaloarchaeota and Micrarchaeota ^5^ have larger genome sizes and seem metabolically more flexible ^14–16,18^; yet cultivated Nanohaloarchaeota representatives are nevertheless host-dependent ^16,17^. While recent work has provided more insights into symbiotic interactions characterizing certain representatives of the DPANN ^16,19^, additional model systems remain to be established.

We have recently succeeded in enriching a member of the Micrarchaeota in a community of four different microorganisms ^15^. Here, we report the isolation of the first stable coculture of this Micrarchaeon together with its host, a previously unknown member of the Thermoplasmatales, as well as the isolation of the latter. This allowed us to conduct experiments aiming to understand the interaction of the two organisms and the response of the Thermoplasmatales member to growth in coculture with the Micrarchaeon.

## Results and discussion

### Isolation of the A_DKE/B_DKE co-culture and pure culture of B_DKE

Originally, we referred to the putative symbionts belonging to the Micrarchaeota and Thermoplasmatales as A_DKE and B_DKE, respectively ^15^. Based on the herein purified co-culture, the isolation of the Thermoplasmatales host the reconstruction of genome sequences of both organisms, we propose the names ‘*Candidatus* Micrarchaeum harzensis’ *sp. nov.* (N.L. masc./fem. adj. *harzensis*, pertaining to the German region of the Harz Mountains, where the organism was isolated) and ‘*Candidatus* Scheffleriplasma hospitalis’ *gen. nov. sp. nov.* (Schef′fler.i.plas′ma. N.L. gen. masc. n. *scheffleri* of Scheffler, named in honor of the geologist Dr. Horst Scheffler and in recognition of his work on mine geology and commitment to our work; Gr. neut. n. *plasma* something shaped or molded; hos.pi.ta′lis. L. masc. adj. *hospitalis* relating to a guest, hospitable, referring to its ability to serve as a host for *Candidatus* ‘Micrarchaeum harzensis*’*). For brevity we will stick to the abbreviation A_DKE and B_DKE throughout this manuscript.

In order to obtain a better understanding of the interactions between A_DKE and B_DKE we first aimed to obtain a co-culture of the putative symbiont and host organisms, i.e. purify the original enrichment from a *Cuniculiplasma*-related archaeon referred to as C_DKE and a fungus (with *Acidothrix acidophila* as its closest related organism) ^15^ C_DKE represented the minority of the archaea in the enrichment culture and is closely related to an organism reported to have a low pH optimum of 1.0–1.2 ^20^. Hence, it was possible to eliminate C_DKE by transferring the culture in media with a pH of 2.5, which exceeds its optimal pH range, while still supporting the growth of A_DKE and B_DKE. Secondly, the fungus, which was isolated and inferred to be psychrophilic/psychrotolerant (data not shown), was successfully eliminated by incubating the enrichment cultures at 37 °C over three consecutive culture transfers. From the previous microscopy analysis, it was known that A_DKE thrives together with B_DKE in biofilm-like structures ^15^. We discovered that it was possible to enhance the biofilm formation of the host- or helper-organism B_DKE by lowering the pH-value of the medium from 2.5 to 2.0, which led to robust biofilm formation of the co-culture (Supplementary Figure S1). At the same time, we were able to isolate B_DKE by enriching at pH 2.5 for planktonic organisms.

### Genomic potential of A_DKE and B_DKE

DNA of the co-culture containing A_DKE and B_DKE was sequenced using a combination of PacBio and Illumina sequencing. For comparison, Illumina sequencing was also performed on the B_DKE pure culture. The two organisms have circular chromosomes of 1 959 588 base pairs (bp) (B_DKE) and 989 838 bp (A_DKE), and GC contents of 44.4 % and 45.8 %, respectively. The genomes of the pure B_DKE isolate and the strain within the co-culture were 100 % identical, which was important for the later comparative transcriptomic analysis. An analysis of clusters of orthologous groups (COGs) revealed that A_DKE contains more proteins with unknown function (29 %; 300 putative proteins without an arCOG-assignment) relative to the overall number of genes compared to B_DKE (20 %; 419 putative proteins without an arCOG-assignment). The complete genome of A_DKE confirmed earlier findings ^15^, such as an extremely limited set of genes coding for proteins involved in central carbon metabolism. We could only detect one gene encoding a putative enzyme of the pentose phosphate pathway and two genes for putative enzymes of a glycolysis or gluconeogenesis pathways (Supplementary Table S1). However, we did identify a putative set of genes coding for enzymes for the conversion of glucose to glycerate, which together comprise four of the seven reactions of the non-phosphorylative Entner-Doudoroff pathway (Supplementary Table S1). The A_DKE genome also contains genes encoding enzymes for the conversion of pyruvate to acetyl-CoA, though we could not identify candidate proteins for reactions leading from glycerate to pyruvate. In agreement with our previous study, proteins for almost all steps of the tricarboxylic acid cycle (TCA) were detected in A_DKE, and may, through the production of NADH, fuel an electron transport chain and the generation of a proton gradient for ATP synthase-based ATP production. Gene clusters encoding a full NADH dehydrogenase and an ATP synthase complex were discovered in the A_DKE genome. Moreover, we identified genes encoding one subunit of the cytochrome bc1 complex and two subunits of the cytochrome c oxidase (Supplementary Table S1). Although the organism might have the ability to generate energy and produce reducing equivalents, it will be highly dependent on building blocks acquired either from the environment (or culture medium) or from the partner organism B_DKE. For instance, A_DKE has major gaps in various biosynthesis pathways including for amino acids; the few steps encoded by A_DKE comprise the production of aspartate from oxaloacetate, glutamate from α-ketoglutarate and phenylalanine from phenylpyruvate. Phenylpyruvate could be produced from tyrosine, which was taken up from the medium by the co-culture (see below). Other amino acid biosynthesis pathways could not be detected. Furthermore, genes encoding known amino acid transporters seem to be absent ^8,10^ and DNA, RNA, and lipid biosynthesis pathways are incomplete (see below). Consequently, A_DKE may acquire certain metabolites or building blocks directly from its partner B_DKE through cell-cell interactions as seen in the *Ignococcus hospitalis/Nanoarchaeum equitans* system ^21,22^. In turn, the dependency of A_DKE on growth in a biofilm (see above) may be due to the need to establish cellular contact with its host.

### Impact of growth in co-culture on the B_DKE transcriptome

To get further insights into the effect of the symbiont on its host, we compared gene expression levels of B_DKE with and without co-cultivation with A_DKE under otherwise identical growth conditions. In particular, we compared three pure cultures with four co-cultures and filtered for differentially expressed genes with p-values lower than 0.05 and log2fold changes higher or lower than 2 or −2. This analysis revealed 15 genes that were differentially expressed based on these criteria (Table 1).

**Table 1:**
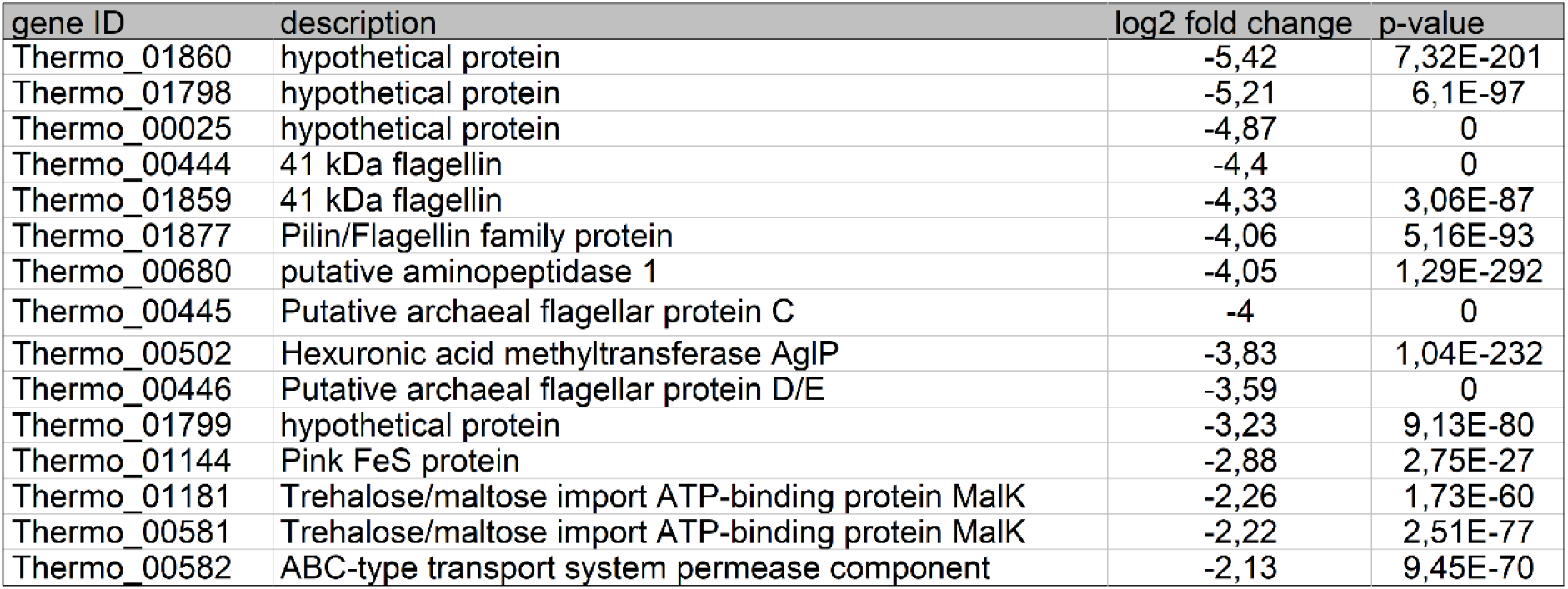
Differentially expressed genes of B_DKE in pure culture in comparison to B_DKE in co-culture with A_DKE. The table indicates the gene ID, the description of the expressed protein, the log2-fold change, and the p-value.

| gene ID | description | log2 fold change | p-value |
| --- | --- | --- | --- |
| Thermo_01860 | hypothetical protein | -5,42 | 7,32E-201 |
| Thermo_01798 | hypothetical protein | -5,21 | 6,1E-97 |
| Thermo_00025 | hypothetical protein | -4,87 | 0 |
| Thermo_00444 | 41 kDa flagellin | -4,4 | 0 |
| Thermo_01859 | 41 kDa flagellin | -4,33 | 3,06E-87 |
| Thermo_01877 | Pilin/Flagellin family protein | -4,06 | 5,16E-93 |
| Thermo_00680 | putative aminopeptidase 1 | -4,05 | 1,29E-292 |
| Thermo_00445 | Putative archaeal flagellar protein C | -4 | 0 |
| Thermo_00502 | Hexuronic acid methyltransferase AgIP | -3,83 | 1,04E-232 |
| Thermo_00446 | Putative archaeal flagellar protein D/E | -3,59 | 0 |
| Thermo_01799 | hypothetical protein | -3,23 | 9,13E-80 |
| Thermo_01144 | Pink FeS protein | -2,88 | 2,75E-27 |
| Thermo_01181 | Trehalose/maltose import ATP-binding protein MalK | -2,26 | 1,73E-60 |
| Thermo_00581 | Trehalose/maltose import ATP-binding protein MalK | -2,22 | 2,51E-77 |
| Thermo_00582 | ABC-type transport system permease component | -2,13 | 9,45E-70 |

All of those 15 genes were downregulated in the host B_DKE in the co-culture compared to the pure culture of B_DKE and comprise five genes of or associated with the B_DKE archaellum complex^23^. This suggests a decreased cellular motility of the host in the co-culture, which is in line with the observed tendency of B_DKE to form a biofilm in the presence of A_DKE. The closest relative of B_DKE, *Cuniculiplasma divulgatum*, does not contain any flagella-related genes^24^, whereas the related ‘G-Plasmas’ are described to contain the full *fla*-operon (*flaBCDEFGHIJ*)^25^. Furthermore, the gene for the hexuronic acid methyltransferase AglP, a component of the protein glycosylation was downregulated and may indicate an alteration of the glycosylation pattern of B_DKE, as well as influence cell-cell-interaction in the presence of A_DKE. It may also change the exopolysaccharide (EPS) matrix of B_DKE, which would explain the observed binding differences of some lectins (see below). Three other downregulated genes code for transport proteins that might be involved in the uptake of carbohydrate molecules. While the effect of the decreased expression level of these transporters is unclear, it may be speculated that it could lead to higher availability of certain metabolites in the medium and support growth of A_DKE. Other downregulated genes encode hypothetical proteins, a putative aminopeptidase, and an iron-sulfur-protein. Thus far, their potential impact on the interaction of B_DKE with A_DKE remains unclear.

### Metabolomic analysis in the presence and absence of A_DKE

Next, we performed a metabolomic analysis to be able to compare metabolites in co-culture to the pure culture and determine whether the presence of A_DKE changes the pattern of depleted and produced organic carbon compounds. The growth of B_DKE in pure culture, estimated on the change in ferrous iron concentration over time, was faster than in the co-culture (Supplementary Figure S2). Therefore, we compared samples with equal ferrous iron concentration (3, 4, and 5 weeks of growth for the pure culture; 5, 6, and 7 weeks of growth for the co-culture) as they indicate similar growth phases. Figure 1 shows a heatmap representing all significantly different metabolites (p < 0.01) present in the samples at the various time points. While the majority of the detected compounds changed simultaneously in the pure- and the co-culture throughout the growth phases, some metabolites showed different patterns.

**Figure 1:**
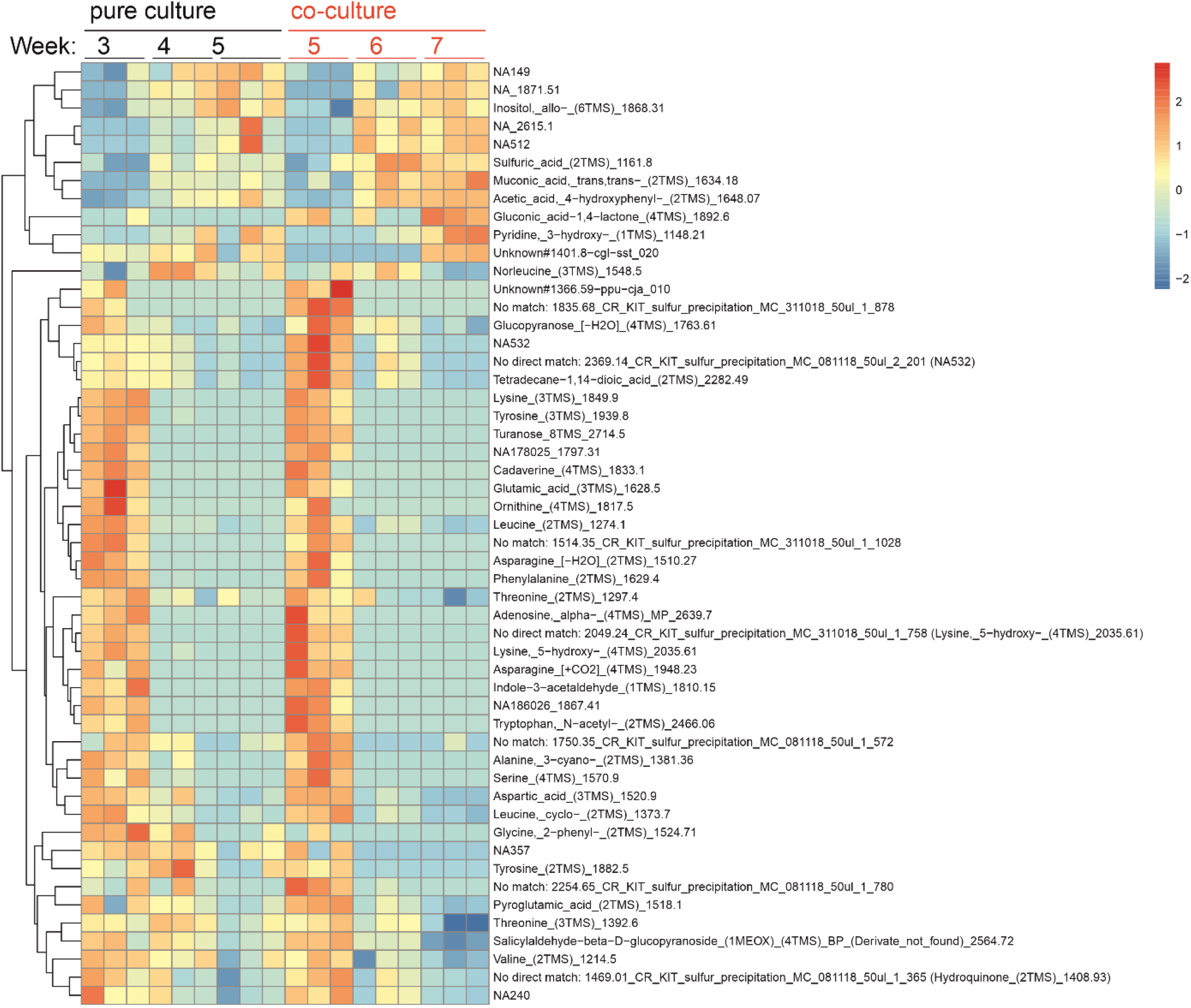
Heatmap showing significantly different metabolite levels (p < 0.01) between corresponding growth phases of pure culture (B_DKE) and co-culture (A_DKE/B_DKE). Normalization was done using a z-score, and significance was calculated by a two-tailed t-test.

Tyrosine levels were found to decrease faster in the co-culture, whereas 4-hydroxyphenyl acetic acid, an intermediate of the tyrosine degradation pathway, increased. Hence, tyrosine degradation seems to be accelerated in the co-culture. Both A_DKE and B_DKE encode enzymes catalyzing the conversion of tyrosine to 4-hydroxyphenylpyuvate (Supplementary table S1) and the presence of the Micrarchaeon might be the reason for faster tyrosine depletion. Note, the specific genes for the carboxylase and dehydrogenase reaction from 4-hydroxyphenylpyruvate to 4-hydroxyphenyl acetic acid are currently unknown. While 2-phenylglycine, a degradation product of phenylalanine that enters the ketoadipate pathway, decreased, muconic acid, an intermediate of that pathway, increased in the co-culture. Still, we could neither identify genes involved in the degradation of 2-phenylglycine nor for a complete ß-ketoadipate pathway in A_DKE or B_DKE (Supplementary Table S1). Moreover, the ß-ketoadipate pathway operates under oxic conditions and the organisms were cultivated in the absence of oxygen. Hence, so far, we cannot explain the consumption and production of 2-phenylglycine and muconic acid, respectively. The analyses also revealed increased levels of gluconic acid in the co-culture, which may be a product of sugar-degradation, for instance from the biofilm EPS matrix ^26^. Genomic information indicates that B_DKE is able to degrade glucose into glucono-1,5-lactone (KO: K18124), which can spontaneously be converted into gluconic acid (Supplementary Table S1). Both organisms possess the enzymes to convert gluconic acid to glycerate, as already discussed above.

Overall, the analysis reveals that the pattern of metabolites does not seem to deviate between isolated B_DKE and co-culture and that the kinetics of consumption show minor differences towards faster consumption of some compounds in the co-culture. Hence, either both organisms employ similar pathways and compounds or, perhaps more likely, A_DKE predominantly uses metabolites provided by B_DKE. The latter assumption is in agreement with the sparsity of transporters encoded in the A_DKE genome and indications for cell-cell interactions among the two organisms.

### Membrane lipids of A_DKE and B_DKE and lipid biosynthetic pathways

An analysis of the intact polar lipids (IPLs) of co-cultures of A_DKE and B_DKE revealed the membrane archaeal isoprenoidal glycerol dibiphytanyl glycerol (GDGT) with zero cyclopentane rings (i.e. GDGT-0, also known as caldarchaeol) as the main lipid, making up to 97 % of the total intact polar lipids, together with a minor amount of archaeol (2,3-di-O-phytanyl glycerol diether) (Supplementary Table S2). Comparison of the results to the pure B_DKE-culture revealed no differences in the relative abundance of the archaeal IPLs, suggesting that the Micrarchaeon A_DKE has an identical membrane lipid composition as its host B_DKE.

Archaeal membrane lipids are formed by isoprenoid side chains linked through ether bonds to glycerol-1-phosphate (G1P, synthesized by the G1P-dehydrogenase^27^) either as a bilayer of diethers (archaeols) or a monolayer of tetraethers (i.e. GDGTs). The isoprenoid building blocks are synthesized by one of the four variants of the archaeal mevalonate (MVA) pathway, which differ with regard to the enzymes mediating the last three enzymatic steps (see ^28^ for a review). Figure 2 shows an overview of the different MVA pathways known. The isoprenoid C20 units are linked to the G1P backbone through ether bonds by the geranylgeranylglyceryl diphosphate (GGGP) synthase and (S)-2,3-di-O-geranylgeranylglyceryl diphosphate (DGGGP) synthase.

**Figure 2:**
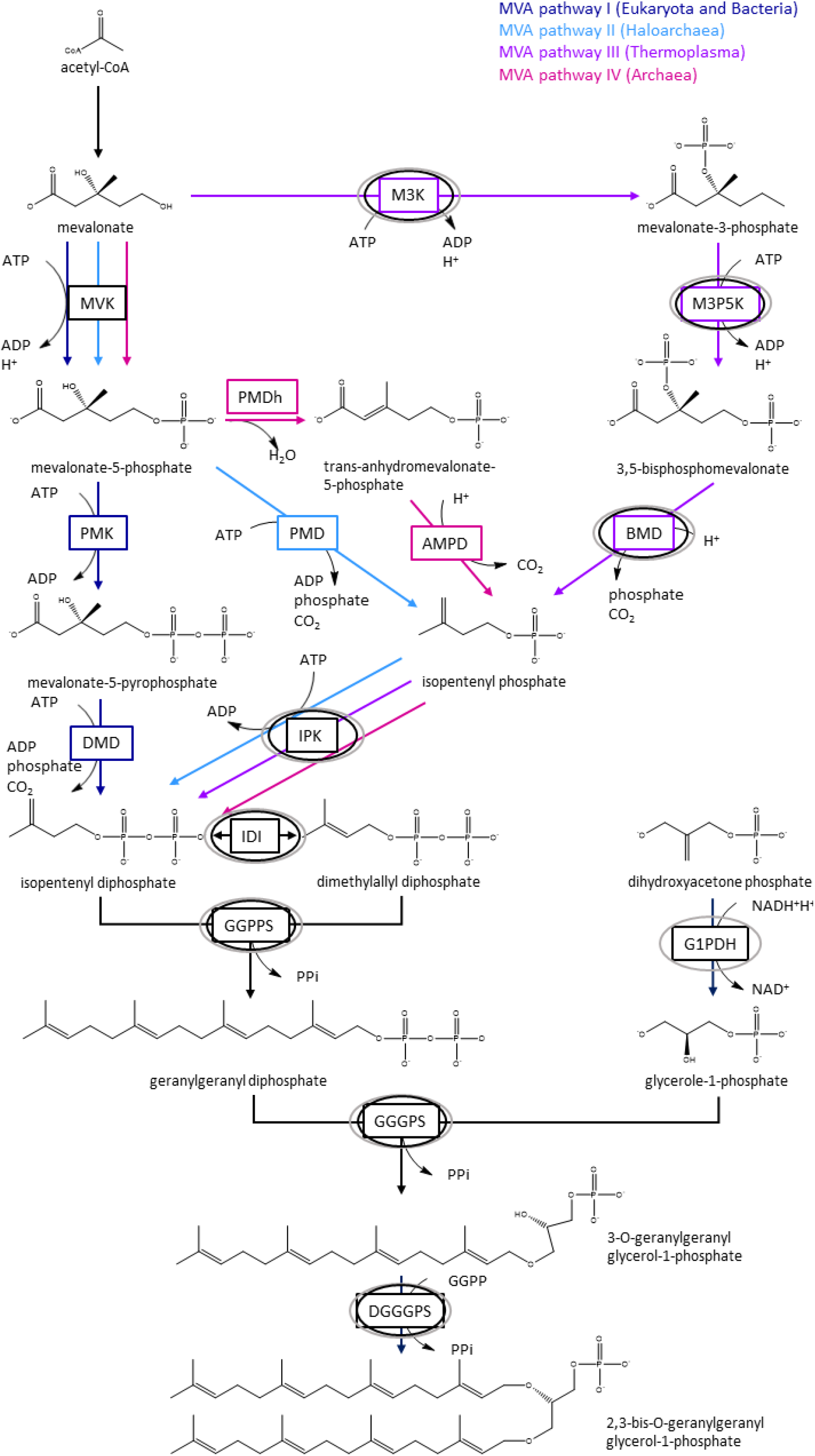
Schematic overview of mevalonate pathways and lipid metabolism. The different pathways are indicated with dark blue (MVA pathway I), light blue (MVA pathway II), violet (MVA pathway III) and magenta arrows (MVA pathway IV), respectively. Names of enzymes are boxed. Enzymes expressed in A_DKE and B_DKE are indicated with black and grey circles. Abbreviations are AMPD: anhydromevalonate phosphate decarboxylase; BMD: bisphosphomevalonate decarboxylase; DGGGPS: 2,3-bis-O-geranylgeranyl glycerol-1-phosphate synthase; DMD: diphosphomevalonate decarboxylase; GGGPS: 3-O-geranylgeranyl glycerol-1-phosphate synthase; GGPP: geranylgeranyl diphosphate; GGPPS: geranylgeranyl diphosphate synthase; G1PDH: glycerol-1-phosphate dehydrogenase; IDI: isopentenyl-diphosphate-delta-isomerase; IPK: isopentenyl phosphate kinase; MVK: mevalonate kinase; M3K: mevalonate 3-kinase; M3P5K: mevalonate 3-phosphate 5-kinase; PMD: phosphomevalonate decarboxylase; PMK: phosphomevalonate kinase.

The genome data (Supplementary Tables S3-S5) revealed that B_DKE uses variant-III of the MVA pathways first described in *Thermoplasma acidophilum* ^29^. In particular, B_DKE encodes the three key enzymes mevalonate-3-kinase (arCOG02937), mevalonate-3-phosphate-5-kinase (COG02074), and mevalonate-3,5-bisphosphate-decarboxylase (arCOG02937), characterizing this pathway, while it lacks genes for a canonical mevalonate kinase (Supplementary Tables S3-S4) similar to other members of the acidophilic Thermoplasmatales (Vinokur *et al.*, 2016). Prenyltransferases found in the genome are farnesyl diphosphate synthase and geranyl-geranyl diphosphate synthase. Also, a G1P-DH (i.e. glycerol-1-phosphate dehydrogenase) could be identified. Genes encoding enzymes for the ether bond formation (i.e. GGGP and DGGGP synthase) and saturation of isoprenoids (i.e. geranylgeranyl reductases) are also encoded in the B_DKE genome (see Supplementary Table S5). In contrast, A_DKE has an incomplete mevalonate and archaeal lipid pathway (Supplementary Tables S3-S4). In particular, while an ancestor of A_DKE and some other Micrarchaeota are likely to have acquired three key genes of the variant-III mevalonate pathway from Thermoplasmatales archaea (i.e. mevalonate-3-kinase (arCOG02937), mevalonate-3-phosphate-5-kinase (COG02074), and mevalonate-3,5-bisphosphate-decarboxylase (arCOG02937)) (Figure 3), A_DKE lacks a homolog of the hydroxymethylglutaryl-CoA reductase (Supplementary Table S3). Note, its genome does not provide any evidence for the presence of another variant of the mevalonate pathway (Supplementary Table S3). Furthermore, A_DKE does not appear to encode a G1PDH. One of the encoded geranylgeranyl reductase homologs of A_DKE, likely involved in lipid biosynthesis, also seems to be acquired by horizontal gene transfer (HGT) from Thermoplasmatales (arCOG00570) (Figure 3).

**Figure 3.**
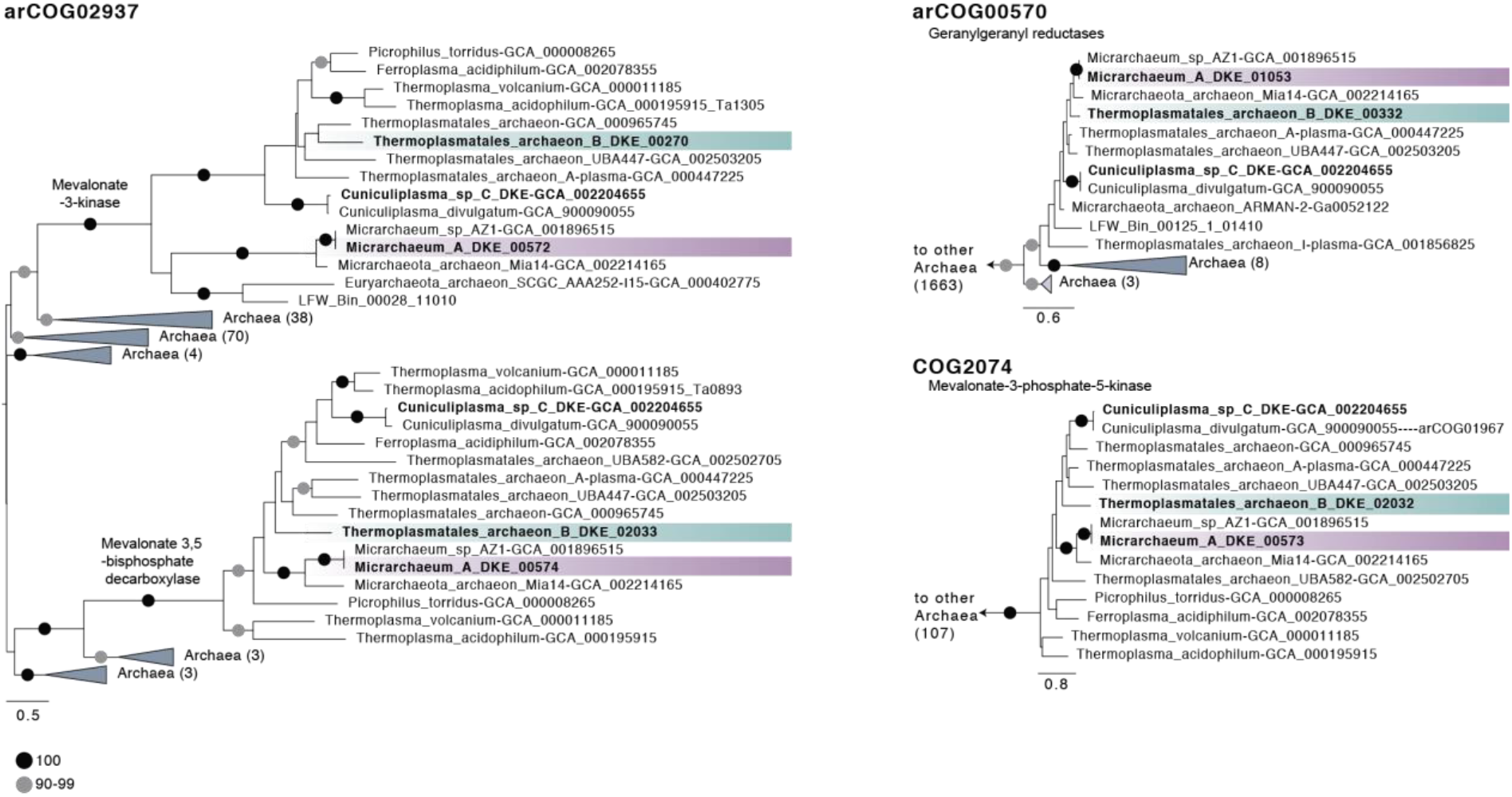
Schematic trees of three protein families with indications for gene transfers between acidophilic Micrarchaeota and Thermoplasmatales. The Maximum likelihood phylogenetic trees shown here included our archaeal backbone dataset (Supplementary Table S6) and were inferred for arCOG02937 (149 sequences with 304 amino acids), arCOG00570 (1877 sequences with 132 amino acids) and COG2074 (arCOG01968 and arCOG01967, 121 sequences with 171 amino acids) with the LG+C10+F+R model with an ultrafast bootstrap approximation run with 1000 replicates. Please note, that the inclusions of bacterial and eukaryotic homologs did not change the interpretation of our findings. Note that arCOG02937 and arCOG02074 represent the key enzymes of the Type-III mevalonate pathway characteristic of Thermoplasmatales. Only bootstrap support values above 90 are shown as indicated in the panel.

Together with our experimental data, the presence of an incomplete variant-III mevalonate and lipid biosynthesis pathways in A_DKE, indicates that this organism depends on lipids or precursors thereof from its host, similar to what has been previously described in the DPANN archaeon *N. equitans* ^31^ and likely other DPANN members such as *Nanohaloarchaeum antarcticus* ^16^.

### Biofilm composition of pure and co-cultures

As the isolation experiments and our previous results point towards the importance of extracellular polymeric substances (EPS) for successful cultivation of A_DKE, we next investigated the composition of the EPS matrix in the co-culture as compared to the pure culture of B_DKE. To this end the glycoconjugates were analyzed with fluorescently labeled lectins, and the signals were correlated to the individual cell type by CARD-FISH analysis (Figure 4). Lectins are complex proteins, which bind specifically to carbohydrate structures ^32^. In this study, 70 different fluorescently labeled lectins, which represent all commercially available lectins available so far (Supplementary Table S7), were used to analyze the EPS in pure and co-cultures.

**Figure 4:**
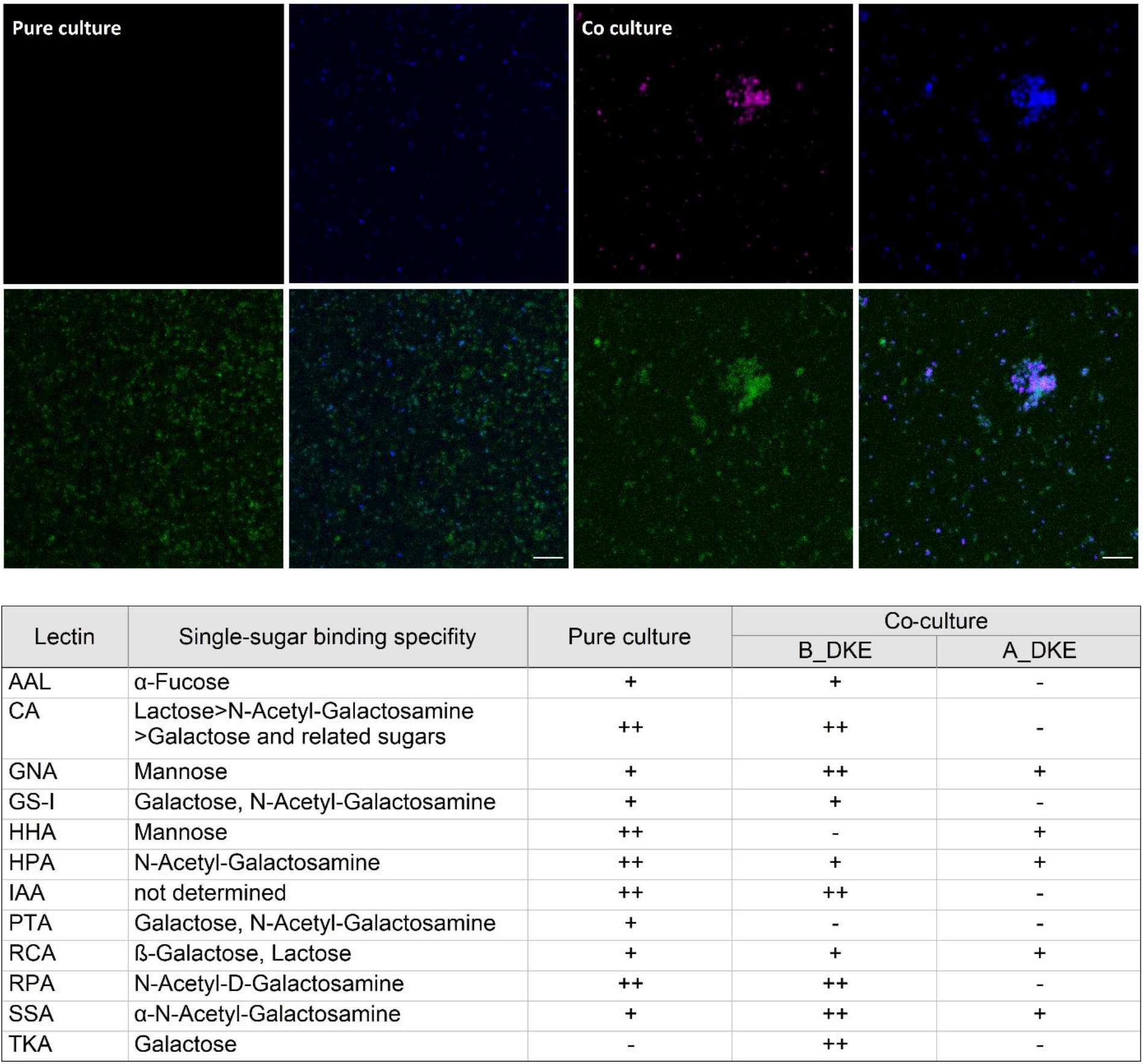
Results of lectin staining of co-culture of A_DKE and B_DKE and B_DKE pure culture. Upper part shows an example of the microscopic analysis of the pure culture (left) and co-culture (right). The images show the results for staining with lectin CA (co-culture) and HHA (pure culture). Color allocation: B_DKE was stained with the general Archaea probe Arch915 (blue) which does not stain A_DKE. A_DKE was stained using the Micrarchaeota-specific ARMAN980 probe (red). Lectin staining is shown in green. Scale bars represent 10 μm. The table shows binding lectins, their carbohydrate-binding specificity, and the strength of the signal in pure and co-culture (with + as weak binding, ++ as binding, and - as no binding).

Among the tested lectins, only those specific to galactose- and mannose-related conjugates bound the extracellular matrix of the co-culture and of B_DKE cells in pure culture. Notably, lectins, IAA, HHA, and PTA seemed to discriminate between the extracellular matrix of B_DKE in the presence or absence of A_DKE, suggesting a potential influence of A_DKE on the matrix chemistry or composition. Lectin staining of A_DKE cells was weak and only possible with some of the lectins that bound to B_DKE. This may reflect the inability of A_DKE to build carbohydrate polymers or the production of a sort of exotic polymer for which we did not have a lectin. It suggests that the detected signals on A_DKE are likely due to growth within the biofilm matrix of B_DKE.

Overall, the results indicate that B_DKE displays galactose and mannose on its cell surface and that these carbohydrates are also components of the co-culture EPS matrix. This is corroborated by the presence of transcriptionally expressed genes for metabolic pathways leading to UDP-glucose, UDP-galactose, GDP-mannose, UDP-N-acetylgalactosamine, and N-acetylglucosamine in the genome of B_DKE (see Supplementary Table S8 and Supplementary Figure S3).

### Evidence for direct cell-cell interactions between A_DKE and B_DKE

Due to the pleomorphic morphology and great variability in cell size of members of the Thermoplasmatales including B_DKE, it was previously challenging to clearly distinguish symbiont and host cells on electron micrographs. Recently, ^33^ has shown that A_DKE cells are characterized by the presence of an S-layer that can be observed on electron micrographs of Platinum-Carbon shadowed samples. Hence, we conducted an electron microscopic study to test whether the two organisms indeed physically interact as indicated by our various analyses (see above). Electron micrographs revealed the attachment of several A_DKE to B_DKE cells, suggesting direct cell-cell interactions between these organisms, as was previously shown for *N. equitans* and *I. hospitalis* ^21,22^ (Figure 5). However, we also observed a large number of A_DKE cells that were not in contact with their potential host organism, which is in agreement with observations from microscopic images of CARD-FISH stained cultures. While we cannot exclude that this is (to some degree) a result of sample preparation, it is possible that growth in the biofilm enables a more dynamic interaction between A_DKE and B_DKE than observed for *N. equitans* and *I. hospitalis* ^21,22^, as the risk of detaching from the host is mitigated by growth within the biofilm matrix. Moreover, A_DKE has a larger genome and in turn greater metabolic flexibility than *N. equitans* ^34^ and may in turn be less dependent on permanent attachment to host cells. Finally, we detected several unattached A_DKE cells in the process of cell division suggesting that A_DKE can store a sufficient amount of building blocks to divide without being in direct cell-cell contact with B_DKE.

**Figure 5:**
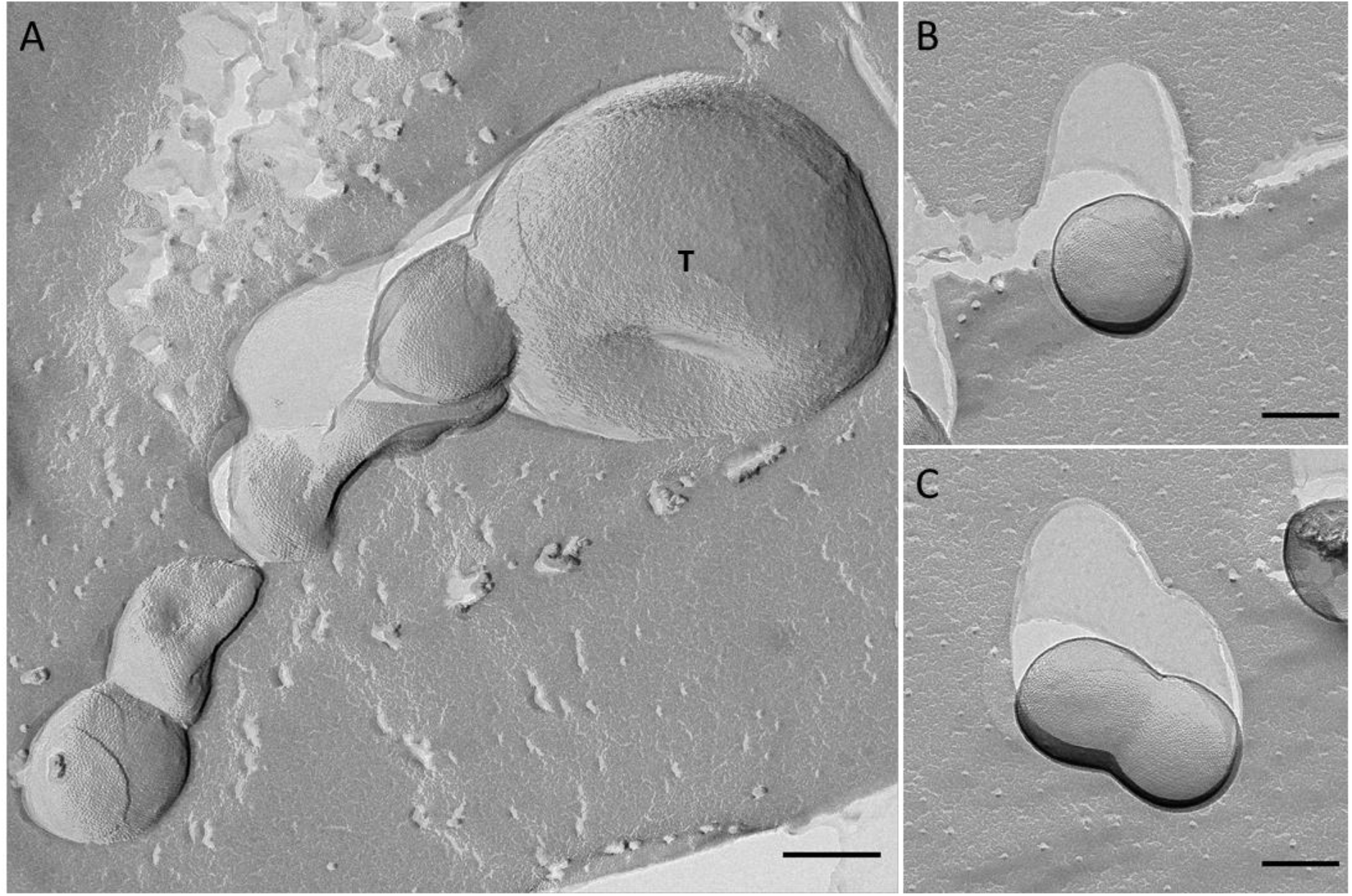
Electron micrographs of freeze-etched, Platinum-Carbon shadowed co-culture cells, containing ‘*Candidatus* Scheffleriplasma hospitalis’ B_DKE and ‘*Candidatus* Micrarchaeum harzensis’ A_DKE. A_DKE cells, displaying an S-Layer on their surface, were observed in physical contact with B_DKE cells (tagged with T) (A), as well as free living (B) and undergoing cell division (C). Scale bars equal 200 nm.

## Conclusion

Previous attempts to purify members of the Micrarchaeota with their respective hosts yielded in relatively diverse enrichments or led to the disappearance of the symbiont after some time of incubation ^14,18,19^. Here we used physiology informed strategies for deselecting against additional community members and show that selection for biofilm formation of the B_DKE host population was crucial for obtaining a stable co-culture. The detailed characterization of our co-culture in comparison with pure host cultures using for instance transcriptomic, metabolomic and lipid analyses indicate a specific regulatory response of B_DKE as a consequence of growth with A_DKE. These characterizations also suggest a dynamic physical interaction of the symbiont and host organisms that is likely crucial for the uptake of various metabolites and building blocks for among others membrane formation. The close cell-cell interactions between acidophilic Micrarchaeota and Thermoplasmatales may also provide a route for horizontal gene transfer among these DPANN symbionts and their hosts (Figure 4). The robust growth of our culture and the possibility to upscale the growth experiments to a fermenter scale provides the basis for prospective work aiming to unravel further insights into the details of the symbiosis between the acidophilic Micrarchaeota A_DKE and its Thermoplasmatales host. On a broader scale, it will be an important step to compare cell-cell interactions underlying this system to the various addition symbioses involving cultivated DPANN archaea and their hosts and establish unique and common characteristics ^12–14,16,17^.

## Materials and Methods

### Culturing conditions

All cultures were cultivated under anoxic conditions in a modified Picrophilus medium at a pH of 2.0 or 2.5 and at 22 °C as previously described ^15^. Transfers of the cultures were conducted with 20 % of the pre-culture. The growth phase was assessed by following the reduction of ferric iron via the ferrozine assay ^35^. Furthermore, the cultures were monitored regarding their activity via CARD-FISH (see below). The cultures typically reached the exponential growth phase after incubation for 2-8 weeks.

### CARD-FISH and Lectin staining

Samples were fixed for 1 h in 4 % formaldehyde, washed twice in phosphate-buffered saline (PBS), and stored at −20 °C in 50:50 PBS/ethanol for CARD-FISH analysis or at 4 °C in 100 % PBS for lectin staining. The fixed cells were hybridized and further processed as described previously ^15,36^. Labeling was conducted using HRP-labelled 16S rDNA probes TH1187 (Thermoplasmatales, GTA CTG ACC TGC CGT CGA C, 20 % formamide; ^37^) and ARM980 (ARMAN, GCC GTC GCT TCT GGT AAT, 30 % formamide; ^38^). For standard CARD-FISH staining, Alexa546 and Alexa488 fluorophores were used, and counterstaining was conducted with DAPI. CARD-FISH staining for lectin analysis was conducted using Alexa546 and Alexa647 fluorophores; see below for more details. Slides were visualized on a Leica DM 5500B microscope (objective lens 100×: HCX PL FLUOTAR, 1.4, oil immersion and objective lens 64x: HCX PL APO; eyepiece 10x: HC PLAN s (25) M), and images were taken with a Leica DFC 360 FX CCD camera and the corresponding Leica LAS AF software.

Lectin staining was conducted according to a protocol of Neu and Kuhlicke (2017). The positive lectin results were compared with CARD-FISH staining of the same slides, using a protocol of Bennke *et al.* (2013). CARD-FISH staining was performed as described above with the following modifications: 1) the low melting agarose and all ethanol washing steps were omitted, as these would negatively affect the lectin staining ^40^; 2) cells were not counterstained with DAPI. After fixation and staining via CARD-FISH, the samples were dried at 37 °C, and 100 μl lectin solution (0.1 mg/mL) was added and incubated for 30 min at room temperature in the dark. After washing with PBS solution, the slides were dried at 37 °C and mounted in the same embedding buffer as used for CARD-FISH. Please refer to Supplementary Table S7 for an overview of lectins used. Imaging was conducted as described above.

### DNA/RNA isolation and quantitative PCR analysis

Isolation of DNA for quantitative (qPCR) analysis was conducted with the Invisorb Spin Forensic Kit following the manufacturer’s instructions (Invitek, Berlin). DNA for Illumina and PacBio sequencing was isolated as described by Lo *et al.* (2017). RNA isolation and library preparation was conducted by IMGM laboratories GmbH using the RNeasy Micro Kit (Qiagen, Hilden) and TruSeq® Stranded total RNA LT kit according to the manufacturer’s instructions (illumina, Berlin). All sequencing analyses were conducted with samples of exponentially growing pure host cultures as well as symbiont-host co-cultures. The ratio of the Micrarchaeon A_DKE to the Thermoplasmatales archaeon B_DKE was calculated via qPCR using standard curves, as described in Krause *et al.* (2017). The primer sequences are listed in Supplementary Table S9.

### Metagenome analysis

A SMRTbell™ template library was prepared according to the instructions from PacificBiosciences, Menlo Park, CA, USA, following the Procedure & Checklist – Greater Than 10 kb Template Preparation. Briefly, for preparation of 15 kB libraries, DNA was end-repaired and ligated overnight to hairpin adapters applying components from the DNA/Polymerase Binding Kit P6 from Pacific Biosciences, Menlo Park, CA, USA. Reactions were carried out according to the manufacturer’s instructions. BluePippin™ Size-Selection to greater than 4 kb was performed according to the manufacturer’s instructions (Sage Science, Beverly, MA, USA). Conditions for annealing of sequencing primers and binding of polymerase to purified SMRTbell™ template were assessed with the Calculator in RS Remote, PacificBiosciences, Menlo Park, CA, USA. One SMRT cell was sequenced on the PacBio RSII (PacificBiosciences, Menlo Park, CA, USA), taking one 240-min movie. Libraries for sequencing on the Illumina platform were prepared to apply the Nextera XT DNA Library Preparation Kit (Illumina, San Diego, USA) with modifications according to Kishony *et al.* ^42^ and sequenced on Illumina NextSeq™ 500 (Illumina, San Diego, USA).

Genome assembly was performed by applying the RS_HGAP_Assembly.3 protocol included in the SMRT Portal version 2.3.0. The assembly revealed two major contigs. Potentially misassembled artificial contigs with low coverage and included in other replicons were removed from the assembly. Redundancies at the ends of the two major contigs allowed them to be circularized. Replicons were adjusted to *smc* (chromosome partition protein Smc) as the first gene. Error-correction was performed by mapping of the Illumina short reads onto finished genomes using the Burrows-Wheeler Aligner bwa 0.6.2 in paired-end mode using default settings ^43^ with subsequent variant and consensus calling using VarScan 2.3.6 (Parameters: mpileup2cns --min-coverage 10 --min-reads2 6 --min-avg-qual 20 --min-var-freq 0.8 --min-freq-for-hom 0.75 --p-value 0.01 --strand-filter 1 --variants 1 --output-vcf 1) ^44^. Automated genome annotation was carried out using Prokka 1.8 ^45^.

### Genome annotations

For further functional annotation, the protein files from the two complete genomes as well as 22 Micrarchaeota and 11 Thermoplasmatales reference genomes (Supplementary Table S6) were compared against several databases, including the Archaeal Clusters of Orthologous Genes (arCOGs ^46^; version from 2018), the KO profiles from the KEGG Automatic Annotation Server (KAAS ^47^; downloaded April 2019), the Pfam database (^48^ Release 31.0), the TIGRFAM database (^49^ Release 15.0), the Carbohydrate-Active enZymes (CAZy) database (^50^ downloaded from dbCAN2 in September 2019), the Transporter Classification Database (TCDB ^51^; downloaded in November 2018), the hydrogenase database (HydDB ^52^; downloaded in November 2018) and NCBI_nr (downloaded in November 2018). Hmmsearch v3.1b298 was used to read HMM profiles of the ArCOG, PFAM, TIGRFAM and CAZyme databases and search against a protein database (settings: hmmsearch <hmmfile> <seqdb> −E 1e-4 ^53^). The best hit for each protein was selected based on the highest e-value and bitscore by using a custom script (hmmsearchTable, available at https://zenodo.org/record/3839790 ^7^). BlastP was used with TCBD, HydDB and NCBI_nr as input databases and the protein sequences as query (settings: −evalue 1e-20 −outfmt 6). Additionally, all proteins were scanned for protein domains using InterProScan (v5.29-68.0; settings: --iprlookup –goterms ^54^). For InterProScan we report multiple hits corresponding to the individual domains of a protein using a custom script (parse_IPRdomains_vs2_GO_2.py) (Supplementary Tables S3 and S4). All custom scripts are available at https://zenodo.org/record/3839790 ^7^.

### Phylogenetic analyses

We performed phylogenetic analyses of membrane lipid biosynthetic proteins, whenever homologs of relevant arCOGs were present in both A_DKE and B_DKE to assess the extent of horizontal gene transfer (HGT) affecting these proteins (Supplementary Table S5). In particular, we extracted homologs of corresponding arCOGs for all of these proteins from A_DKE, B_DKE, a reference set of 566 archaeal genomes (archaea-only analysis) as well as from an additional of 3020 bacterial and 100 eukaryotic genomes (universal analysis) (Supplementary Table S6). The reference genomes were annotated as described above. For the archaea-only analysis, the individual homologs for each protein family were aligned using MAFFT L-INS-i v7.407 (settings: --reorder ^55^), trimmed with BMGE v1.12 (settings: −t AA −m BLOSUM30 −h 0.55 ^56^). Subsequently, phylogenetic trees were inferred using IQ-TREE (v1.6.10, settings: −m LG+C10+F+R −wbtl −bb 1000 −bnni ^57^). For the universal analysis, MAFFT L-INS-i v7.407 and MAFFT v7.407 were used to align protein families with Less/equal (≤) or more (>) than 1000 homologs, respectively. BMGE v1.12 was used for trimming all alignments (settings: −t AA −m BLOSUM30 −h 0.55 ^56^) and phylogenetic trees were inferred using IQ-TREE (v1.6.10, settings: −m LG+C10+F+R −wbtl −bb 1000 −bnni ^57^). Due to the large number of sequences affiliating with arCOG00570 when including bacterial and eukaryotic homologs (i.e. 13381), sequences for this protein family were aligned using MAFFT v7.407, trimmed with TrimAL (v1.2rev59, settings: −gappyout ^58^). Sequences with => 90 % gaps were removed using a custom script (faa_drop.py) and used for a phylogenetic analysis with FastTree (v2.1.10, settings: −lg −gamma).

### Transcriptomic analysis

Sequencing of RNA samples of three replicates of the pure culture and four replicates of the co-culture was performed on an Illumina NextSeq® 500 NGS system using 2 × 75 bp paired-end read chemistry. All samples were taken from cultures in exponential growth phase. Transcriptomic analyses were performed using Kallist v0.45.0 ^59^ and compared to the reference completed genomes (see above). Differential expression was assessed by using the R package DESeq2 (1.24.0; Love *et al.*, 2014). Log2fold change shrinkage for normalization was calculated by the ashr program ^61^.

### Lipid analyses

For the analysis of membrane lipids, cells were derived from 20 mL pure and co-culture in the exponential growth phase. Samples were taken for monitoring of cells via CARD-FISH prior to filtering (Supplementary Figure S4). Cells were filtered onto 0.3 μm pore size 47 mm diameter glass fiber filters (GF75, Advantec MFS, Inc, CA, USA). Total lipids were extracted from the freeze-dried glass-fiber filters using a modified Bligh and Dyer method ^62^, as described earlier ^63^. The extracts were dried under nitrogen and split into two aliquots, one left untreated and another hydrolyzed with 1.5 M HCl in methanol by reflux at 130 °C for 2 h to remove the headgroups from the archaeal intact polar lipids (IPL) and release the core lipids (CLs). The pH was adjusted to 7 by adding 2 M KOH/MeOH (1:1 v/v) and, after the addition of water to a final 1:1 (v/v) ratio of H_2_O-MeOH, extracted three times with dichloromethane (DCM). The DCM fractions were collected and dried over sodium sulfate. The dried samples were dissolved in hexane–2-propanol (99:1, vol/vol) and filtered over a 0.45-μm polytetrafluoroethylene filter. The extracts after acid-hydrolysis contained the IPL-derived CLs plus CLs, while the non-hydrolyzed extracts consisted only of the CLs. Extracts were analyzed by UHPLC–atmospheric pressure chemical ionization (APCI) MS for archaeal CLs, including archaeol (diether, C_20_ isoprenoid chains) and glycerol dialkyl glycerol tetraether (GDGTs, tetraether, C_40_ side chain), according to Hopmans *et al.* (2016), with some modifications. Briefly, the analysis was performed on an Agilent 1260 UHPLC coupled to a 6130 quadrupole MSD in selected ion monitoring (SIM) mode. This allowed the detection of GDGTs with 0 to 4 cyclopentane moieties, crenarchaeol as well as archaeol. The separation was achieved on two UHPLC silica columns (BEH HILIC columns, 2.1 × 150 mm, 1.7 μm; Waters) in series, fitted with a 2.1 × 5-mm pre-column of the same material (Waters) and maintained at 30 °C. Archaeal CLs were eluted isocratically for 10 min with 10 % B, followed by a linear gradient to 18 % B in 20 min, then a linear gradient to 100 % B in 20 min, where A is hexane and B is hexane:isopropanol (9:1). The flow rate was 0.2 mL/min. Total run time was 61 min with a 20 min re-equilibration. Source settings were identical to Schouten *et al.* (2007). The typical injection volume was 10 μl of a 1 mg/mL solution (weighted dried Bligh and Dyer extract dissolved in hexane:isopropanol (99:1, v/v ratio). The m/z values of the protonated molecules of archaeol and GDGTs were monitored. GDGTs were quantified by adding a C_46_ GTGT internal standard ^66^. A response factor derived from an archaeol:GDGT-0 standard (1:1) was used to correct for the difference in ionization between archaeol and GDGTs.

The Bligh and Dyer extract (non-hydrolyzed) and the acid-hydrolyzed Bligh and Dyer extract were also analyzed using ultra-high-performance liquid chromatography coupled to positive ion atmospheric pressure chemical ionization/Time-of-Flight mass spectrometry (UHPLC-APCI/ToFMS) on an Agilent 1290 Infinity II UHPLC, equipped with an automatic injector, coupled to a 6230 Agilent TOF MS and Mass Hunter software. This additional analysis was performed to detect other archaeal lipids that were not included in the SIM method on the 6130 quadrupole MSD mentioned above. Separation of the archaeal lipids was achieved according to Hopmans *et al.* (2016) with some modifications using two silica BEH HILIC columns in series (2.1 × 150 mm, 1.7 μm; Waters) at a temperature of 25 °C. The injection volume was 10 μL. Compounds were isocratically eluted with 90 % A and 10 % B for the first 10 min, followed by a gradient to 18 % B in 15 min, a gradient to 30 % B in 25 min, and a linear gradient to 100 % B in 30 min. A = hexane and B = hexane/isopropanol (9:1, v/v) and the flow rate was 0.2 mL/min. The conditions for the APCI source were identical to Schouten *et al.* (2007) and Hopmans *et al.* (2016). Also, the fragmentor was set at 300 V. The ToFMS was operated in extended dynamic range mode (2 GHz) with a scan rate of 2 Hz. We assessed archaeal lipid distributions by monitoring m/z 600 to 1400. Archaeal lipids were identified by searching within 10 ppm mass accuracy for relevant [M+H]+ signals.

### Metabolome analysis

A pure culture of B_DKE and a co-culture of A_DKE and B_DKE were inoculated as described above for 42 (pure culture) or 49 days (co-culture) until all available ferric iron was reduced and stationary phase was reached. Experiments were performed in triplicates. For metabolomic examination, 1 mL culture was sampled and stored at −80 °C until further analyses. Samples were taken every 7 days, along with samples for ferrous iron quantification to estimate growth, and samples for DNA extraction and CARD-FISH for further detection of A_DKE and B_DKE. Due to the low pH of the culture medium, 500 μl samples were amended by inducing sulfur precipitation through the addition of a spatula tip of CaCO_3_ to each sample, mixing for 5 min at 2000 rpm at room temperature. This treatment also led to cell lysis so that the analysis included also intracellular metabolites. After centrifugation for 5 min at 14000 rpm at room temperature, 50 μl of the supernatant was transferred to glass vials and dried under vacuum at 4 °C. Dried samples were stored at −80 °C until further analysis.

Online metabolite derivatization was performed using a Gerstel MPS2 autosampler (Muehlheim, Germany). Dried metabolites were dissolved in 15 μl of 2 % methoxyamine hydrochloride in pyridine at 40 °C under shaking. After 90 min, an equal volume of N-Methyl-N-(trimethylsilyl)trifluoroacetamide (MSTFA) was added and held for 30 min at 40 °C. One μl of the sample was injected into an SSL injector at 270 °C in splitless mode. Gas chromatography/mass spectrometry (GC/MS) analysis was performed using an Agilent 7890A GC equipped with a 30-m DB-35MS # 5-m Duraguard capillary column. Helium was used as carrier gas at a flow rate of 1.0 mL/min. The GC oven temperature was held at 100 °C for 2 min and increased to 300 °C at 10 K/min. After 3 min, the temperature was increased to 325 °C. The GC was connected to an Agilent 5975C inert XL MSD, operating under electron ionization at 70 eV. The MS source was held at 230 °C and the quadrupole at 150 °C. The total run time of one sample was 60 min. All GC/MS chromatograms were processed by using the Metabolite Detector software ^67^.

### Structural analysis by electron microscopy

For freeze-etching, both a pure culture of B_DKE and a co-culture of A_DKE and B_DKE were concentrated by centrifugation (3,000 x g). The concentrated cell pellet (1.5 μL) was applied onto a gold carrier, frozen in liquid nitrogen, and transferred into a freeze-etching device (CFE-50, Cressington, Watford, UK; p < 10^−5^ mbar). At T=176 K, samples were fractured using a cold knife (T=90 K); after sublimation of about 400 nm of surface water, the samples were shadowed with Platinum-Carbon at an angle of 45 degrees (1.5 nm), and an additional layer of pure Carbon (about 15 nm; both by electron-beam evaporation). Replicas were cleaned for 15 hours on 70 % H2SO4, washed three times on bidistilled water, taken up on 700 mesh (hex) grids and air-dried. For electron microscopy analysis at 200 kV, a transmission electron microscope JEM-2100F (JEOL GmbH, Freising, Germany), equipped with a F416 CMOS camera (TVIPS, Gauting, Germany) under control of SerialEM v. 3.8 ^68^ was used.

## Accession of data

The genome sequences including annotations have been deposited at NCBI Genbank under Accession Numbers CP060530 and CP060531. Raw reads of transcriptomic data are available under SRA files SRX8933312-SRX8933318. All raw files phylogenetic trees can be found in a repository (doi: 10.5281/zenodo.4725436).

## Acknowledgements

The authors thank Carola Berg for excellent technical assistance.

