## Supplemental Information for "Unraveling the critical growth factors for stable cultivation of (nano-sized) Micrarchaeota"

Supplementary information

**Suppl. Table S1**

Genes of A_DKE with a function in central metabolic pathways with gene ID and annotation. Annotations according to Suppl. Table S4 and S5.

**Suppl. Table S2**

Relative abundance of lipids in co-culture (B_DKE+A_DKE) and pure culture (B_DKE). Abbreviations are CL: core lipids, IPL: intact polar lipids.

**Suppl. Table S3**

Table showing the annotation of protein files of A_DKE and Micrarchaeota reference genomes across different databases.

**Suppl. Table S4**

Table showing the annotation of protein files of B_DKE and Thermoplasmatales reference genomes across different databases.

**Suppl. Table S5**

Table listing genes used for phylogenetic analyses.

**Suppl. Table S6**

Table listing reference genomes used for phylogenetic analysis.

**Suppl. Table S7**

Table of all used lectins with Carbohydrate binding specificity and linkage type.

**Suppl. Table S8**

Genes in A_DKE and B_DKE genome with function in glycosylation and synthesis of carbohydrate precursors for EPS matrix. Listed are the gene ID, gene product, arCOG/TIGR numbers and e-values, respectively, as well as TPM expression values in co- and pure culture. Annotations were done with KEGG Annotation Server and compared to results of Suppl. Table S4-S5.

**Suppl. Table S9**

Primers used for calculation of the ratio of A_DKE to B_DKE via qPCR.

**Suppl. Figure S1**

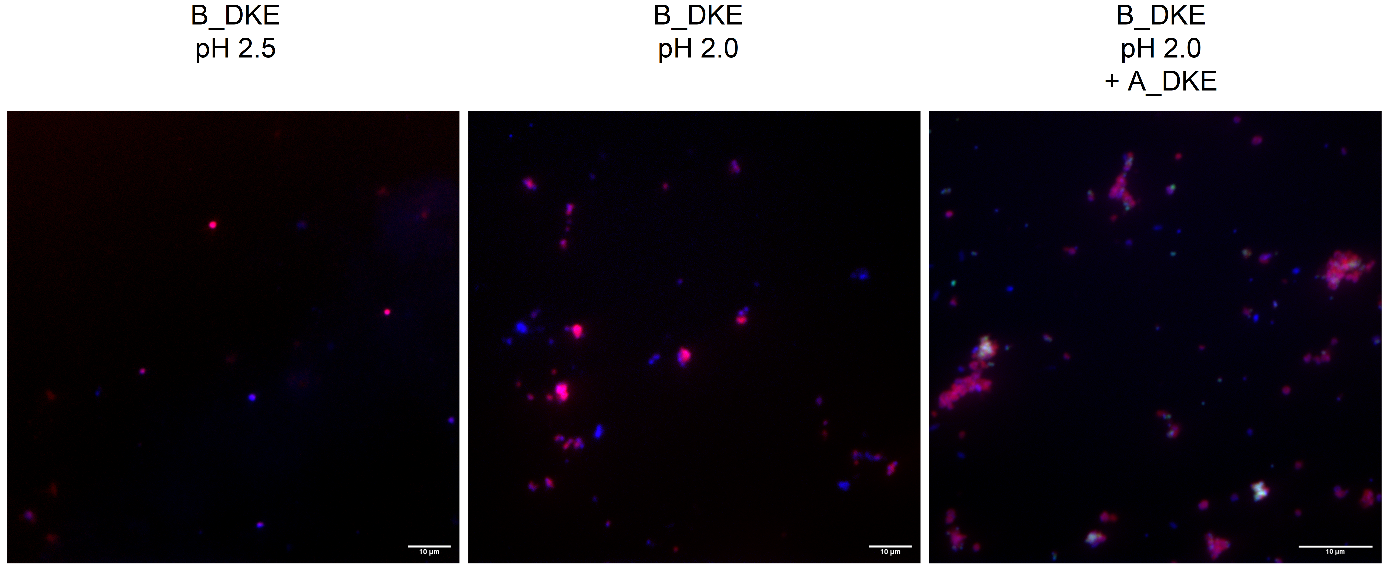

Suppl. Figure S2: CARD-FISH pictures depicting growth morphology of B_DKE cells under different conditions. Microscopic pictures showing (from left to right) B_DKE cultivated at pH 2.5, cultivated at pH 2.0 and co-cultivated with A_DKE at pH 2.0. Samples were taken at similar growth phases. Cells are stained as follows: B_DKE in red (TH1178 probe), A_DKE in green (ARM980 probe) and all cells in blue (DAPI).

**Suppl. Figure S2**

Fe2+ [mM]

| time [weeks] | mean pure culture | mean co-culture |
| --- | --- | --- |
| 0 | 6.16136 | 4.20364 |
| 2 | 11.8917 | 6.52265 |
| 3 | 20.2627 | 6.85874 |
| 4 | 31.4148 | 10.4549 |
| 4.85714 | 38.0838 | 16.1846 |
| 6 | 42.7595 | 29.2085 |
| 7 |  | 40.2261 |

Suppl. Figure S3: Fe^2+^ concentration during growth curve of co-culture of A_DKE and B_DKE and pure culture of B_DKE for metabolic analysis. Data shown, are mean values of a triplicate, respectively.

**Suppl. Figure S3**

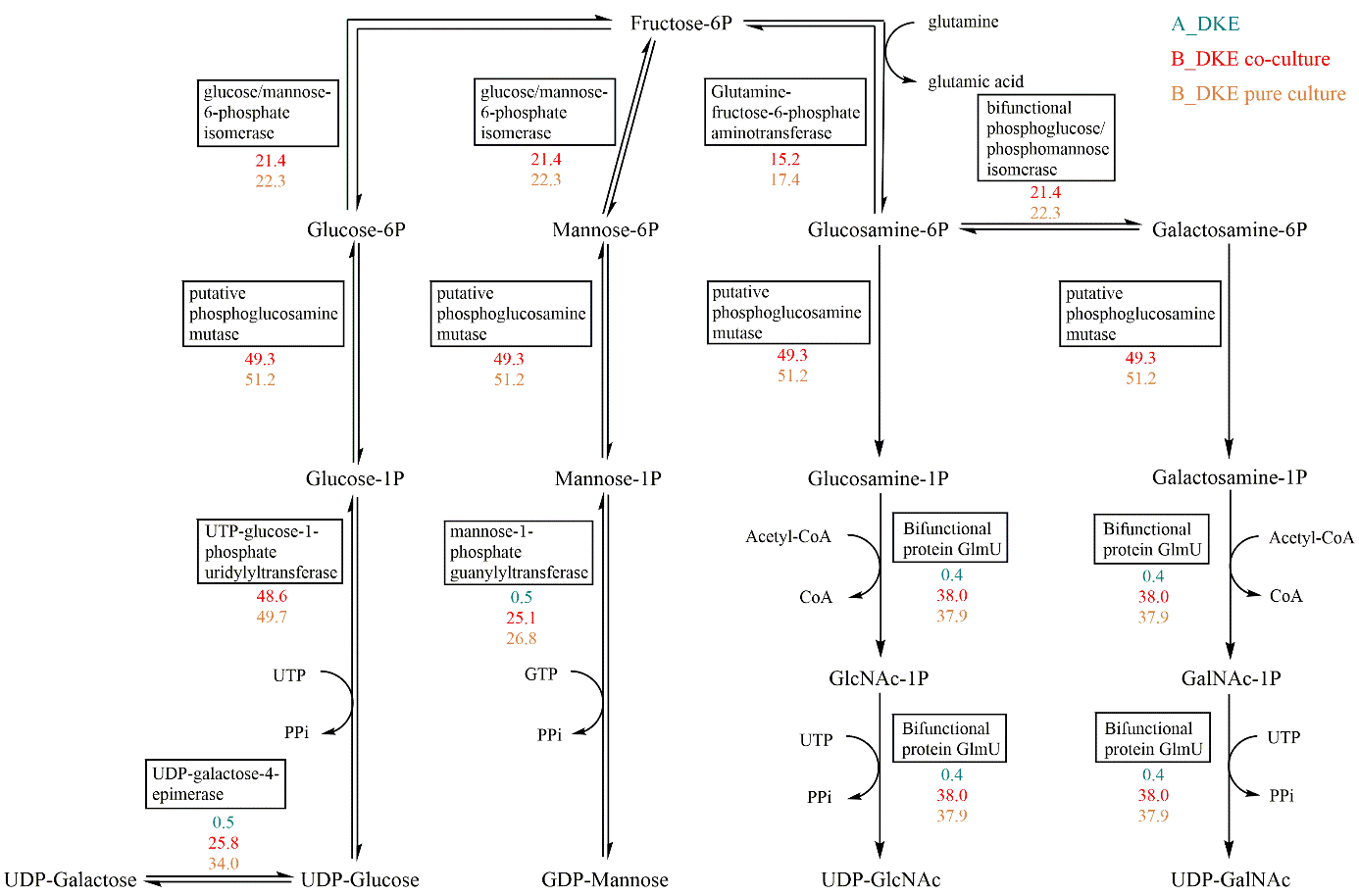

Suppl. Figure S4: Depiction of the metabolic pathways of EPS precursors from fructose-6-phophate. The TPM values of the responsible enzymes are listed in green (A_DKE), red (B_DKE in co-culture) and orange (B_DKE in pure culture). Abbreviations are; UTP: uridine 5’-triphosphate; UDP: uridine 5’-diphosphate; GlcNAc: N-acetyl-glucosamine; GalNAc: N-acetyl-galactosamine.

**Suppl. Figure S4**

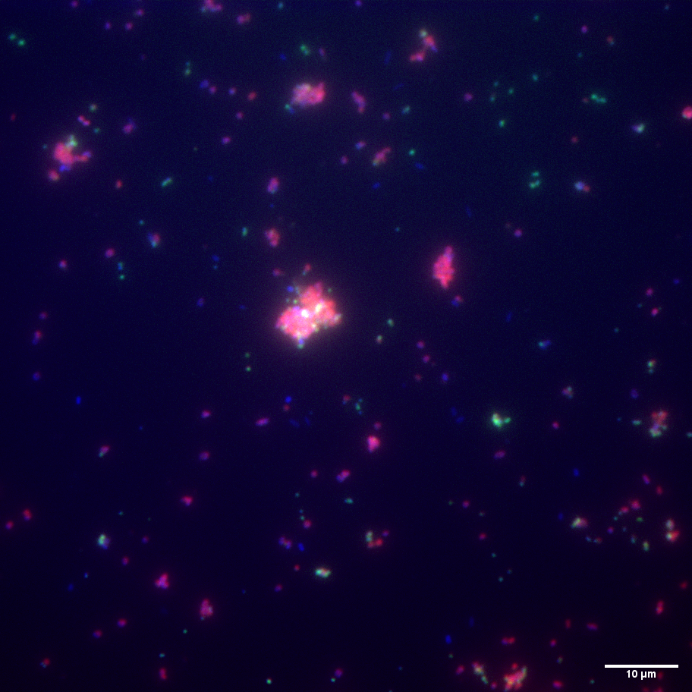

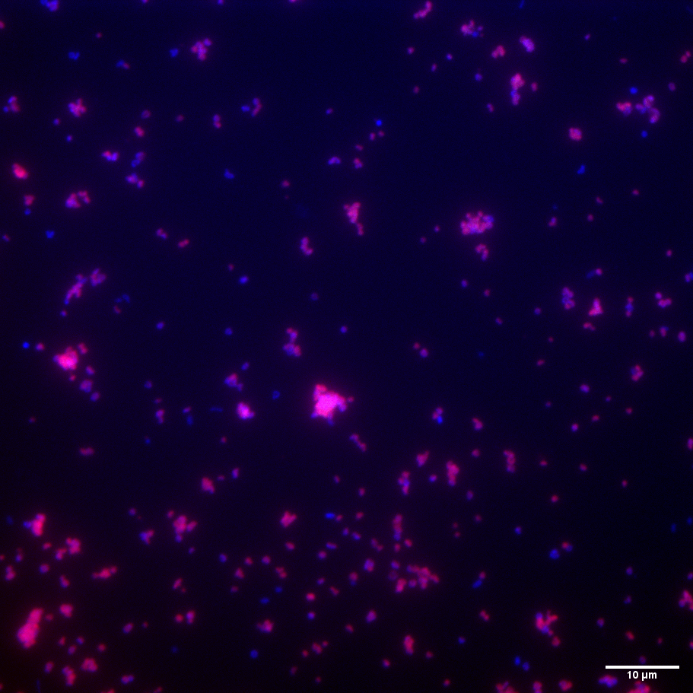

Suppl. Figure S1: CARD-FISH pictures of cultures used for lipid analysis. Left picture shows an overlay of the used co-culture containing A_DKE and B_DKE, right pictures an overlay of the pure culture of B_DKE. Red: TH1178 probe staining B_DKE, green: ARM980 probe staining A_DKE, blue: DAPI as counter staining.
